# Feminization of social play behavior depends on microglia

**DOI:** 10.1101/2024.08.19.608675

**Authors:** Jonathan W VanRyzin, Ashley E Marquardt, Margaret M McCarthy

## Abstract

Many sex differences in brain and behavior are established developmentally by the opposing processes of feminization and masculinization, which manifest following differential steroid hormone exposure in early life. The cellular mechanisms underlying masculinization are well-documented, a result of the fact that it is steroid-mediated and can be easily induced in newborn female rodents via exogenous steroid treatment. However, the study of feminization of particular brain regions has largely been relegated to being “not masculinization” given the absence of an identified initiating trigger. As a result, the mechanisms of this key developmental process remain elusive. Here we describe a novel role for microglia, the brain’s innate immune cell, in the feminization of the medial amygdala and a complex social behavior, juvenile play. In the developing amygdala, microglia promote proliferation of astrocytes equally in both sexes, with no apparent effect on rates of cell division, but support cell survival selectively in females through the trophic actions of Tumor Necrosis Factor α (TNFα). We demonstrate that disrupting TNFα signaling, either by depleting microglia or inhibiting the associated signaling pathways, prevents the feminization of astrocyte density and increases juvenile play levels to that seen in males. This data, combined with our previous finding that male-like patterns of astrocyte density are sculpted by developmental microglial phagocytosis, reveals that sexual differentiation of the medial amygdala involves opposing tensions between active masculinization and active feminization, both of which require microglia but are achieved via distinct processes.

**Significance Statement:** The cellular mechanisms by which sex differences in the brain arise provide insight into the cellular basis of behavior. Most mechanistic studies have focused on the process whereby regions of the male brain are differentiated from the female in response to elevated gonadal steroid in development due to the tractability of inducing masculinization by blocking steroid action in males or providing exogenous steroids to newborn females. As such, feminization is usually defined as “not masculinized”. Here, we demonstrate the active feminization of astrocyte density in a brain region modulating complex social behavior, rough-and-tumble play in juveniles. These findings indicate that lower levels of playfulness in females is an actively regulated process as opposed to simply being a lack of masculinization.

## INTRODUCTION

Many sex differences in the brain are established developmentally by the process of sexual differentiation. Occurring during a critical period of perinatal development in rodents, two opposing processes act to achieve sexual differentiation of the brain: masculinization, which is induced by androgens and aromatized estrogens, and feminization, which occurs in the absence of elevated steroid hormones. Many regions of the brain that are subject to sexual differentiation are embedded within the social behavior network (Newman 1999; Goodson 2005), including the medial preoptic nucleus, ventromedial hypothalamus, bed nucleus of the stria terminalis, anteroperiventricular nucleus, and the medial amygdala. Sex differences in these regions arise from carefully orchestrated organization of developmental processes such as synaptic patterning, cell genesis, and cell death and enduring epigenetic modifications (McCarthy & Arnold, 2011; Forger, 2016; McCarthy et al., 2017; Tsukahara & Morishita, 2020). While the cellular underpinnings of masculinization are well-studied (Zuloaga et al., 2008), the mechanisms underlying feminization of brain and behavior remain poorly understood. This is in large part due to the lack of a definable initiating signal, such as testosterone, as occurs in males.

Microglia, once considered mere sentinels of the brain’s immune system, are now appreciated for their complex suite of abilities that are fundamental to the formation of a functional brain. Microglia coordinate circuit development by promoting and pruning synaptic connectivity (Paolicelli et al., 2011; Schafer et al., 2012; Ji et al., 2013; Squarzoni et al., 2014; Miyamoto et al., 2016), regulating cell proliferation via trophic factor support, and inducing cell death by phagocytosing dead/dying cells or cells targeted for engulfment (Marin-Teva et al., 2004; Sierra et al., 2010; Cunningham et al., 2013; Ueno et al., 2013; Shigemoto-Mogami et al., 2014; Hagemeyer et al., 2017). Microglia are also essential to sexual differentiation, as many sex differences in behavior arise from microglia functions that are induced by the divergent steroid hormonal milieu, such as dendritic spine induction and phagocytosis (Lenz et al., 2013; VanRyzin et al., 2019; Pickett et al., 2023).

The medial amygdala (MeA) is a critical node for processing social information and is one of the brain regions most strongly impacted by hormone-mediated sexual differentiation, with profound importance for adult social and sexual behavior (Choi et al., 2005; Bergan et al., 2014; Unger et al., 2015; Ishii et al., 2017; Li et al., 2017; Lischinsky et al., 2017; Raam & Hong, 2021; Lischinsky et al., 2023). The MeA is also uniquely involved in mediating the sex difference in frequency and intensity of juvenile rough-and-tumble play as a result of androgen-induced masculinization during the perinatal critical period (Meaney et al., 1983; Meaney & McEwen 1986). We have previously discovered that microglia mediate masculinization of the MeA, as microglia phagocytose newborn astrocyte progenitors more frequently in the amygdala of males than females. This results in decreased astrocyte density in the posterodorsal medial amygdala (MePD) of juvenile males compared to females, which is permissive for increased MePD neural activity and higher rates of rough-and-tumble play (VanRyzin et al., 2019).

Here, we expand upon this framework and describe a novel role for microglia in feminization of the MeA and juvenile play. We show that microglia promote cell proliferation in the developing MeA equally in both sexes, but actively support cell survival selectively in females. Treating females with testosterone eliminates the cell survival capabilities of microglia. The microglia-mediated surviving cells are largely astrocytes, and their survival is driven by the trophic actions of Tumor Necrosis Factor α (TNFα), which is more highly expressed by microglia in females. Disrupting TNFα signaling, either by decreasing TNFα via microglia depletion or by inhibiting downstream TNFα signaling cascades, prevents the feminization of newborn cell number and increases later-life juvenile play behavior. Together, these findings demonstrate that the sexual differentiation of the MeA is the result of a multifaceted process whereby active feminization is opposed to active masculinization, with microglial regulation of the astrocyte population as a central and causal mechanism.

## MATERIALS AND METHODS

### Animal studies

Adult male and female Sprague-Dawley rats were purchased from Charles River Laboratories and delivered to the University of Maryland School of Medicine animal facility. Rats were kept on a 12:12 h reverse light/dark cycle with ad libitum food and water and mated in our facility. Pregnant females were checked daily for pups and the day of birth was designated as postnatal day 0 (P0). On P0, pups were sexed and culled to no more than 14 per dam. Both male and female pups were used in these studies and treatment and sex were balanced across litters. All animal procedures were performed in accordance with the regulations of the Institutional Animal Care and Use Committee at the University of Maryland School of Medicine.

### Animal treatments

Animals were treated with the following pharmacological agents and dosages according to the timelines indicated in the figures and text:

BrdU (50 mg/kg; Signa-Aldrich #B5002) dissolved in saline and delivered intraperitoneally at volume of 0.1 mL per day.

EdU (50 mg/kg; Invitrogen #A10044) dissolved in saline and delivered intraperitoneally at a volume of 0.1 mL per day.

Testosterone proprionate (100 µg; Sigma-Aldrich #T1875) dissolved in sesame oil and delivered subcutaneously in a volume of 0.1 mL per day.

Liposomal clodronate (Encapsula Nanosciences #CLD-8909) or empty liposome vehicle delivered via bilateral intra-amygdalar injection (1 µL per hemisphere) each day at the following coordinates with reference to bregma: –0.8 mm A/P, ±3.0 mm M/L, –5.0 mm D/V.

Celastrol (0.5 μg; Tocris #3203) dissolved in saline and delivered via bilateral intracerebroventricular injection (1 µL per hemisphere) each day at the following coordinates with reference to bregma: –1.0 mm A/P, ±1.0 mm M/L, –3.0 mm D.

Intracranial injections were performed under cryoanesthesia using a 23 gauge Hamilton syringe attached to a stereotaxic frame. The time pups were separated from the dam was 15 min to 1 h.

### Immunofluorescence

Rats were anesthetized with Fatal Plus (Vortech Pharmaceuticals) and transcardially perfused with phosphate-buffered saline (PBS; 0.1M, pH 7.4) followed by 4% paraformaldehyde (PFA; 4% in PBS, pH 7.2). Brains were removed and postfixed for 24 hours in 4% PFA at 4°C, then kept in 30% sucrose at 4°C until fully submerged, rapidly frozen and coronally sectioned at a thickness of 45 µm (for developmental studies) or 20 µm (for juvenile studies) on a cryostat (Leica CM2050S) and directly mounted onto slides.

Slides were washed with PBS and blocked with 5% bovine serum albumin (BSA) in PBS + 0.4% Triton X-100 (PBS-T) for 1 h. Slides were incubated in primary antibody solution (2.5% BSA in PBS-T) overnight at RT. The following day, slides were incubated in secondary antibody solution (2.5% BSA in PBS-T) for 2 h, stained with Hoescht 33342 (Thermo Fisher Scientific) or DAPI (Thermo Fisher Scientific), and coverslipped with ProLong Diamond Antifade Mountant (Thermo Fisher Scientific).

BrdU labeling included an additional antigen retrieval step (0.01M sodium citrate, pH 6.0 for 20 min at 99°C) prior to the blocking step to enhance signal detection. Co-labeling with EdU proceeded according to manufacturer’s instructions (Click-iT Plus EdU Kit; Thermo Fisher Scientific; #C10637). The following primary antibodies were used in this study: Rabbit anti-Iba1 (1:1000; Wako #019-19741), mouse anti-BrdU (1:500; BD Biosciences #347580), rabbit anti-GFAP (1:1000, Abcam #7260). Secondary antibodies used in this study included: Alexa Fluor (all 1:500; Thermo Fisher Scientific) donkey anti-rabbit 488 (#A21206), donkey anti-rabbit 594 (#A21207), and donkey anti-mouse 594 (#A21203).

### Stereological cell counting

Unbiased stereological cell counting was performed using StereoInvestigator software (MBF Bioscience) and a Nikon Eclipse E600 microscope equipped with a MBF Bioscience CX9000 camera. Every third section (45 µm thick) throughout the amygdala was used for analysis, for a total of four sections, and the amygdala from both hemispheres was quantified. The boundaries of the amygdala were identified using a neonatal rat atlas as reference (Ashwell and Paxinos, 2008). The optical fractionator method was used to quantify BrdU+ cells and pyknotic bodies at 40x magnification. BrdU+ cells and pyknotic bodies were quantified using a 50 µm x 50 µm counting grid with a 100 µm x 100 µm sampling grid. Optical dissector height was set to 12 µm with a 2 µm guard zone at the top and bottom for all quantifications.

### *In situ* hybridization

In situ hybridization was performed using the RNAscope Multiplex Fluorescent Assay v2 according to manufacturer’s instructions (ACD Bio). In brief, animals were transcardially perfused with PBS followed by 4% PFA, removed, and postfixed for 24 hours in 4% PFA at RT. Brains were transferred to 30% sucrose solution until fully submerged, then rapidly frozen and cut at a thickness of 20 µm on a cryostat (Leica CM2050S) and mounted onto slides. Sections were dehydrated with ascending ethanol and pretreated with hydrogen peroxide followed by protease III. Sections were then processed according to the RNAscope protocol, including probe incubation and fluorescence labeling. For colocalization with Iba1 immunofluoresence, tissue was processed for immunohistochemistry (as described above) immediately following the final washes of the RNAscope protocol. RNAscope probes targeting TNFα (*TNF*; #402671), TNFR1 (*TNFRSF1A*; #408111), TNFR2 (*TNFRSF1B*; #1312151), and Ki67 (*MKI67*; #515801) were used in combination with Opal dyes (Akoya Biosciences) 520, 620, 650.

### Image acquisition

For all experiments, confocal images were acquired using a Nikon A1 equipped with 405, 488, 561, and 647 lasers and a 20x (0.75 NA), 40x (0.95 NA), 60x (1.40 NA) oil-immersion, or 100x (1.45 NA) oil-immersion objective, or a Zeiss LSM 800 equipped with 405, 488, 561, and 640 lasers and a 20x (0.8 NA), or 63x (1.4 NA) oil-immersion objective. Images were deconvolved prior to analysis using Nikon Elements software or Zeiss Zen software.

### Quantification of microglia morphology and TNFα expression

Coronal sections (45 μm thick) were labeled for TNFα transcript expression by RNAscope and subsequently immunolabeled for Iba1 as described above. Imaging was performed using confocal microscopy and images were acquired using a 40x objective and 0.5 μm z-step through the entire tissue thickness. The Surfaces Module within Imaris (Bitplane) was used to generate 3D-reconstructions of individual microglia. Each 3D rendering was then used as a masking boundary to isolate TNFα signal within each microglia. The Surfaces Module was again used to calculate the volume of the microglia specific TNFα signal and normalized to the volume of each microglia to determine relative expression as a percentage of each microglia volume. Microglia morphology was assessed as done previously (Schwarz et al., 2012; VanRyzin et al., 2019).

### Quantification of GFAP+ cell density

Coronal sections (20 μm thick) were immunolabeled for GFAP and imaged via confocal microscopy as described above. Images were acquired using a 20x objective and 1 μm z-steps through the entire tissue thickness, and the resulting maximum intensity projection was used for cell counting using Imaris (Bitplane). The number of GFAP+ cells was normalized to the area of the MePD to account for any volumetric differences between sexes or treatments. GFAP+ cells were included in the analysis if a single nucleus could be identified and was associated with the GFAP signal.

### Quantification of RNAscope colocalization

Coronal sections (45 μm thick) were labeled for TNFR1, TNFR2, and Ki67 transcript expression by RNAscope and imaged via confocal microscopy as described above. Images were acquired using a 20x objective and 1 μm z-steps through the entire tissue thickness, and the resulting maximum intensity projection was used to determine colocalization using Imaris (Bitplane). Transcripts were considered to be colocalized if signals were present within the boundaries of a complete nucleus.

### Juvenile social play testing

Behavior testing was conducted approximately 2 h after the start of the dark phase of the light cycle under red light illumination. On P21, animals were weaned and housed into same-sex, same-treatment sibling pairs. Play behavior testing occurred once per day from P27-P30 as previously described (VanRyzin et al., 2020a). Same-sex, same-treatment non-sibling pairs of animals were placed in an enclosure (24 x 18 x 12 in) with TEK-Fresh cellulose bedding (Harlan Laboratories). Animals were allowed to acclimate to the chamber for 2 min, then were video recorded for behavioral analysis for 10 min. Videos were scored offline by an experimenter blind to treatment condition and sex to determine the number of pounces, pins, and boxing behaviors.

### RNA extraction, cDNA synthesis, and quantitative PCR

RNA was extracted from amygdala tissue punches using the RNeasy Mini Kit (Qiagen #74104) according to the manufacturer’s protocol. Single-stranded cDNA synthesis was performed using the high-capacity cDNA Reverse Transcription Kit (Thermo Fisher Scientific #4368814). Quantitative PCR (qPCR) was conducted using an Applied Biosystems ViiA7 PCR System (Thermo Fisher Scientific). Primers (Integrated DNA Technologies) were designed using Primer3 software, and efficiencies were determined through serial dilution. All samples were run in triplicate and cycle threshold (Ct) values were normalized to the Ct for *Gapdh* (Δ-Ct), and relative expression was determined by the ΔΔ-Ct method (Schmittgen and Livak, 2008). The following primers were used in this study: *TNFα*: FWD: 5’-GTGATCGGTCCCAACAAGGA-3’ REV: 5’-CGCTTGGTGGTTTGCTACGA-3’; *IGF1*: FWD: 5’-ATCATGTCGTCTTCACATCTCTTCTAC-3’ REV: 5’-CTGTGGGCTTGTTGAAGTAAAAGC-3’; *VEGFa*: FWD: 5’-GCCCACGTCGGAGAGCAA-3’ REV: 5’-CTTTGTTCTATCTTTCTTTGGTCTGCAT-3’; *BDNF*: FWD: 5’-GCGGCAGATAAAAAGACTGC-3’ REV: 5’-CAGTTGGCCTTTTGATACCG-3’; *CD11b*: FWD: 5’-GGTGGTGATGTTCAAGCAGAATTT-3’ REV: 5’-GGTATTGCCATCAGCGTCCAT-3’; *TGFβ1*: FWD: 5’-CAATTCCTGGCGTTACCTTGGT-3’ REV: 5’-CCCTGTATTCCGTCTCCTTGGT-3’; *NGF*: FWD: 5’-CCACTCTGAGGTGCATAGCGTAA-3’ REV: 5’-GTGGGCTTCAGGGACAGAGTCT-3’; *GAPDH*: FWD: 5’-TGGTGAAGGTCGGTGTGAACGG-3’ REV: 5’-TCACAAGAGAAGGCAGCCCTGGT-3’.

### Quantification and Statistical Analysis

All quantifications were conducted by experimenters blind to treatment and sex. Statistical analyses were performed using R (R Core Team, 2021; version 4.1.0) or GraphPad Prism (version 10.2.3). All values are shown as the mean ± SEM. Comparisons between two experimental groups were performed using a two-tailed Student’s t test. Data including multiple experimental groups were analyzed using two-way analysis of variance (ANOVA). Bonferroni’s post hoc comparisons were calculated for specific comparisons to determine differences between male and female groups and within-sex effects of treatment. Pearson’s r was used to calculate linear correlation. A p value of < 0.05 was used to determine significance. Additional statistical details of specific experiments can be found in figure legends and in the text.

## RESULTS

### Microglia Depletion Decreases Cell Proliferation in the Developing Amygdala

We previously observed high rates of microglial phagocytosis of newborn cells in the developing amygdala, of which a large majority (∼83%) were destined to become astrocytes (VanRyzin et al., 2019). We therefore reasoned that eliminating microglia would increase the number of astrocytes surviving until the juvenile period. To test that hypothesis, we treated male and female rat pups with liposomal clodronate via intra-amygdalar injection on P1 and P2, which we previously demonstrated reduces Iba1+ cells in the amygdala by ∼85% by P4 (VanRyzin et al., 2019). To mark cells born during the period of microglia depletion, we injected pups with the thymidine analog 5-bromo-2’-deoxyuridine (BrdU) once on P2 and quantified the number of BrdU+ cells on P4 (Figure 1A). Contrary to our prediction, liposomal clodronate treatment significantly and substantially (∼50%) reduced the number of BrdU+ cells in both males and females (Figure 1B, 1C).

**Figure 1.**
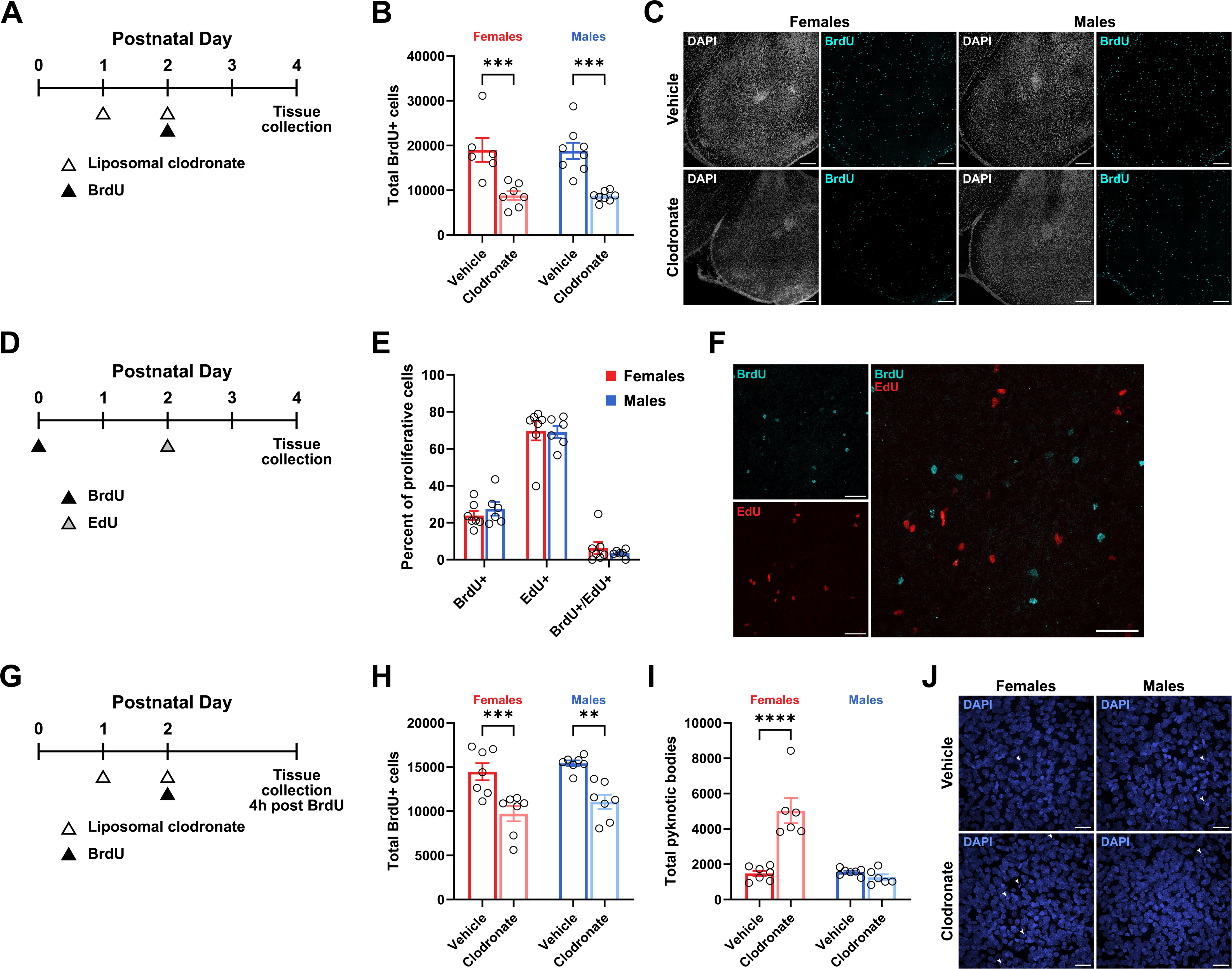
Microglia Depletion Decreases Cell Proliferation in Both Sexes but Increases Cell Death Selectively in Females. (A) Schematic showing the experimental treatments and timeline for (B) and (C). (B) Quantification of the number of BrdU+ cells. Two-way ANOVA main effect of treatment F(1, 25) = 41.26; p < 0.0001. Bonferroni’s post hoc comparisons are shown, n = 6-8 rats per group. (C) Representative images of the amygdala from female (left) and male (right) animals treated with vehicle (top) or clodronate (bottom) immunolabeled for DAPI and BrdU. Scale bars represent 200 µm. (D) Schematic showing the experimental treatments and timeline for (E) – (F). (E) Quantification of the percentage of proliferating cells that immunolabeled as BrdU+, EdU+, and BrdU+/EdU+ in the male and female amygdala. n = 6-7 rats per sex. (F) Representative field of view of the amygdala immunolabeled for BrdU (top left) and EdU (bottom left) and resulting merged image (right). Scale bars represent 50 µm. (G) Schematic showing the experimental treatments and timeline for (H) – (J). (H) Quantification of the number of BrdU+ cells. Two-way ANOVA main effect of treatment F(1, 24) = 33.71; p < 0.0001. Bonferroni’s post hoc comparisons are shown, n = 7 rats per group. (I) Quantification of the number of pyknotic bodies. Two-way ANOVA sex x treatment interaction F(1, 22) = 30.59; p < 0.0001. Bonferroni’s post hoc comparisons are shown, n = 6-7 rats per group. (J) Representative field of view of the amygdala from female (left) and male (right) animals treated with vehicle (top) or clodronate (bottom) labeled for DAPI. White arrowheads indicate pyknotic bodies. Scale bars represent 25 µm. Bars represent the mean ± SEM. Open circles represent individual data points for each animal. **p < 0.01, ***p < 0.001, ****p < 0.0001.

A reduction in the number of newborn cells can be achieved in multiple ways, such as a decrease in the number of cell divisions, a slowing of the rate of proliferation, or an increase in newborn cell death. To rule out the possibility that the decrease in BrdU+ cell number was due to a decrease in the number of cell divisions of highly proliferative progenitor cells, we combined BrdU labeling with a second thymidine analog, 5-Ethynyl-2-deoxyuridine (EdU) and determined the percentage of proliferating cells that were BrdU+/EdU+, indicating successive cell divisions during the limited injection period. Male and female pups were injected with BrdU on P0 followed by EdU on P2 and immunohistochemically assessed for cell proliferation on P4. Approximately 25% of newborn cells in both sexes were born on P0 (i.e. BrdU+ alone; 27.51% ± 3.62% in males; 23.89% ± 2.47% in females) with the majority (∼69%) of newborn cells born on PN2 (i.e EdU+ alone; 68.98% ± 3.26% in males; 69.71% ± 5.16% in females). Interestingly, only a very small percentage (∼5%) of cells born on PN0 divided again on PN2 (i.e. co-labeled BrdU+ and EdU+; 3.51% ± 0.82% in males; 6.39% ± 3.19% in females; Figure 1E), suggesting that the majority of cell proliferation in the amygdala during the early postnatal window is likely the result of a few division cycles from lowly proliferative progenitors.

We next tested the possibility that differences in the total number of proliferative cells may be the source of the observed decrease in newborn cells. Newborn pups P1 and P2 were treated with liposomal clodronate to deplete microglia as before, administered BrdU on P2, and tissue collected for analysis four hours after BrdU injection (Figure 1G). This time course represents approximately two BrdU half-lives and is significantly shorter than the time for a full cell division cycle (∼24 hours: Ponti et al., 2013), thus ensuring that the BrdU+ cells analyzed represent an accurate “snapshot” of cell proliferation. As before, clodronate treatment reduced BrdU+ cell number in both males and females (Figure 1H), confirming that microglia depletion significantly decreases cell proliferation in the developing amygdala.

Finally, to rule out cell death as a contributing factor, we also quantified pyknotic bodies in tissue from these same animals to determine the extent of cell death during the period of microglia depletion. Surprisingly, we found an increase in the number of pyknotic bodies only in females treated with liposomal clodronate, with no effect in males or between vehicle-treated males and females (Figure 1I, 1J). Together, these data suggest that microglia regulate cell proliferation similarly in males and females but influence cell death in a sex-specific manner.

### Microglia Regulate Cell Survival in Females, Feminizing Juvenile Play Behavior

We next sought to determine the influence microglia have on newborn cell survival in the amygdala during the early postnatal window. Toward that goal, we reversed the order of treatment and injected male and female pups with BrdU on P0 to mark newborn cells, followed by microglia depletion with liposomal clodronate on P1 and P2 (Figure 2A). Since BrdU was administered 24 hours prior to liposomal clodronate treatment in this paradigm, typical rates of cell proliferation should be preserved; thus, any observed differences in BrdU-labeled cell count on P4 are likely attributable to the loss of secreted proteins, such as trophic factors, released by microglia. Interestingly, clodronate treatment significantly decreased BrdU+ cell number only in females (Figure 2B), suggesting that microglia-mediated cell survival is biased towards females.

**Figure 2.**
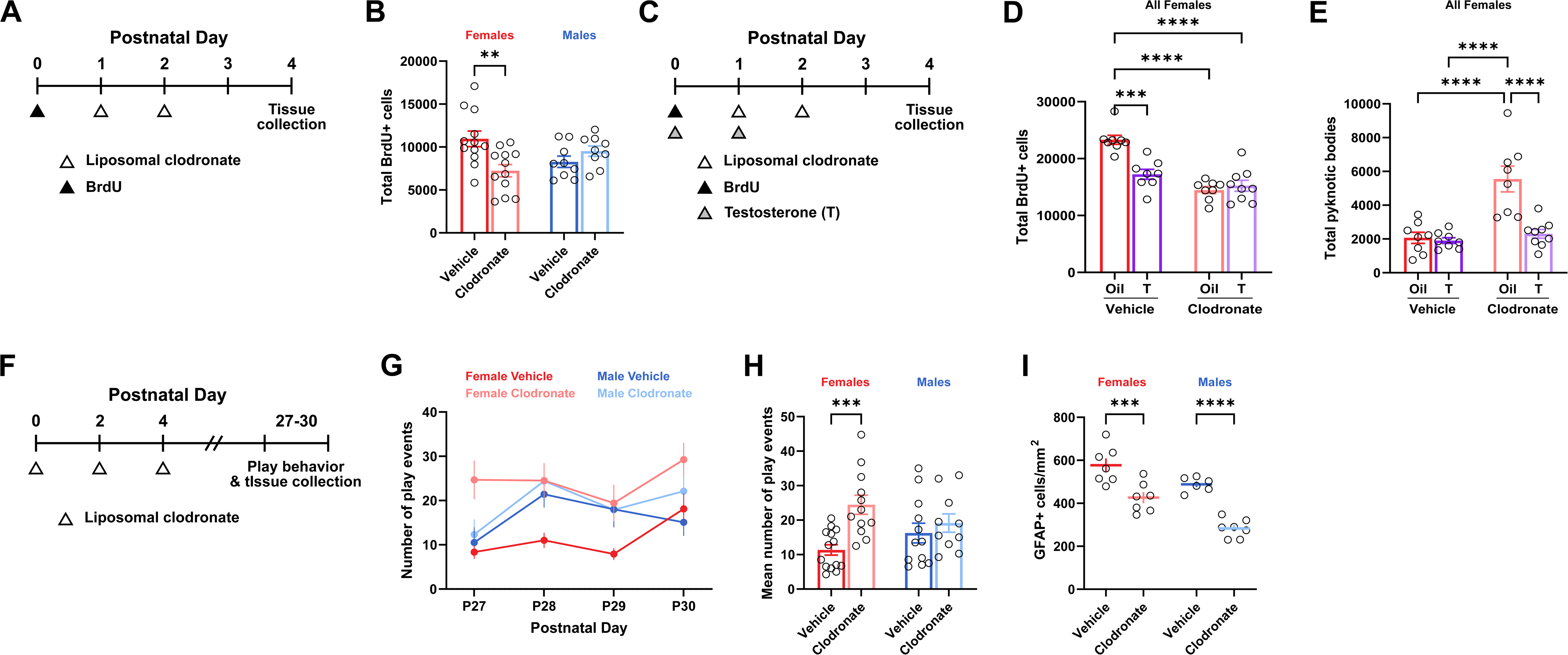
Microglia Depletion Decreases Cell Survival Selectively in Females in a Testosterone-Dependent Manner. (A) Schematic showing the experimental treatments and timeline for (B). (B) Quantification of the number of BrdU+ cells. Two-way ANOVA sex x treatment interaction F(1, 38) = 10.32; p = 0.0027. Bonferroni’s post hoc comparisons are shown, n = 9-12 rats per group. (C) Schematic showing the experimental treatments and timeline for (C) – (D). (D) Quantification of the number of BrdU+ cells. Two-way ANOVA hormone x treatment interaction F(1, 29) = 16.58; p = 0.0003. Bonferroni’s post hoc comparisons are shown, n = 8-9 rats per group. (E) Quantification of the number of pyknotic bodies. Two-way ANOVA hormone x treatment interaction F(1, 29) = 12.60; p = 0.0013. Bonferroni’s post hoc comparisons are shown, n = 8-9 rats per group. (F) Schematic showing the experimental treatments and timeline for (G) – (I). (G) Quantification of the number of play events daily from P27 to P30. Colors indicate sex and treatment. (H) Quantification of the mean number of play events from P27 to P30. Two-way ANOVA sex x treatment interaction F(1, 44) = 4.318; p = 0.0436. Bonferroni’s post hoc comparisons are shown, n = 10-14 rats per group. (I) Quantification of the density of GFAP+ cells in the MePD. Two-way ANOVA main effect of sex F(1, 23) = 23.53, p < 0.0001; main effect of treatment F(1, 23) = 55.51; p < 0.0001. Bonferroni’s post hoc comparisons are shown, n = 6-7 rats per group. Bars and line graphs represent the mean ± SEM. Open circles represent individual data points for each animal. MePD, posterodorsal medial amygdala. **p < 0.01, ***p < 0.001, ****p < 0.0001.

We next reasoned that the effect of microglia depletion on cell survival selectively in females may be due to their lack of androgen exposure typically experienced by males during the critical period for sexual differentiation of the brain. To test this hypothesis, we again treated female pups with BrdU on P0, combined with a masculinizing dose of testosterone on P0 and P1 and liposomal clodronate on P1 and P2 (Figure 2C). We predicted that females masculinized via testosterone treatment would exhibit the result we previously observed in males: no effect of liposomal clodronate on BrdU+ cell count at P4. As expected, testosterone-treated (masculinized) females treated with empty liposomes resembled males, in that they had significantly fewer BrdU+ cells compared to oil-treated (non-masculinized), non-depleted females. However, in females whose microglia were depleted, the number of BrdU+ cells was reduced regardless of masculinization status (Figure 2D). We quantified the number of pyknotic bodies in the same sections and found a significant increase in pyknotic bodies only in control females treated with liposomal clodronate, as seen previously (Figure 1I). Importantly, testosterone– and clodronate-treated females had a similarly low level of pyknotic bodies as compared to control (oil– and vehicle-treated females (Figure 2E). Together, these data demonstrate testosterone masculinizes female brain development by effectively rendering the developmental program “insensitive” to the cell-survival effects of microglia. We presume this process occurs naturally in males in response to their own prenatal androgen production but cannot test this directly as the exposure to androgen in males begins in utero 3-5 days prior to birth.

Next, we sought to determine the functional impact of the female-biased decrease in cell survival in the amygdala. In our previous studies, juvenile playfulness was inversely correlated with newborn cell number during postnatal development in the amygdala: males had fewer newborn cells early in life, and subsequently had lower astrocyte density and correspondingly higher rates of rough-and-tumble play as juveniles (Krebs-Kraft et al., 2010; VanRyzin et al., 2019). Here, we treated male and female pups with liposomal clodronate on P0, P2, and P4 to deplete microglia throughout the early postnatal window and assessed play behavior from P27 – P30 (Figure 2F). Juvenile play significantly increased in females treated with clodronate as neonates compared to control-treated females. There was no effect of neonatal microglia depletion in males (Figure 2G, 2H), likely because microglia depletion also eliminates two other microglia functions that affect later-life play, namely microglia-mediated phagocytosis and microglia-mediated proliferation support, which act to oppose each other during this period of development.

To determine if the impact of microglia depletion on play behavior in females was attributable to alterations in astrocyte density, we quantified astrocytes using the astrocyte-specific protein, glial fibrillary acidic protein (GFAP), via immunohistochemistry in the MePD. Neonatal microglia depletion decreased GFAP+ cell density in the juvenile MePD in both males and females (Figure 2I), congruent with an expected decrease in astrocytes due to the reductions in newborn cell number due to the loss of microglia-mediated proliferation support (Figure 1B). We interpret these data to indicate that play behavior is masculinized in females by the combination of the loss of the pro-proliferative effects and the cell survival effects of microglia, thereby reducing astrocyte number in the amygdala, a process that normally occurs in males in response to androgen-promoted microglia-mediated phagocytosis.

### TNFα and TNFα Receptors on Microglia are Sex-Biased in the Neonatal Amygdala

To identify which microglial factor was responsible for regulating newborn cell survival in the developing amygdala in females, we analyzed the expression of select cytokines and trophic factors known to be produced by microglia and/or regulate cell proliferation and death in the developing brain, including *TNFα*, *TGFβ*, *NGF*, *BDNF*, *IGF1*, and *VEGF* (O’Kusky et al., 2000; Arnett et al., 2001; Butovsky et al., 2006; Pérez-Martín et al., 2010; Shigemoto-Mogami et al., 2014; Kreisel et al., 2019; Okabe et al., 2020). We quantified expression by qPCR in tissue samples of amygdala from P4 animals after liposomal clodronate treatment from P0-2 (Figure 3A). *TNFα* expression was dramatically reduced in microglia-depleted males and females (Figure 3B). The microglial marker *CD11b* was quantified as a measure of depletion efficacy and showed a similarly dramatic reduction (Figure 3C). Levels of *TNFα* and *CD11b* expression significantly correlated in both males and females (Figure 3D), suggesting microglia are a significant and likely primary source of TNFα in the developing brain. Of the other factors assessed, microglia depletion only marginally decreased expression of *TGFβ* in males and females (Figure 3E) and had no effect on the expression of *NGF* (Figure 3F), *BDNF* (Figure 3G), *IGF1* (Figure 3H), and *VEGF* (Figure 3I) in either sex.

**Figure 3.**
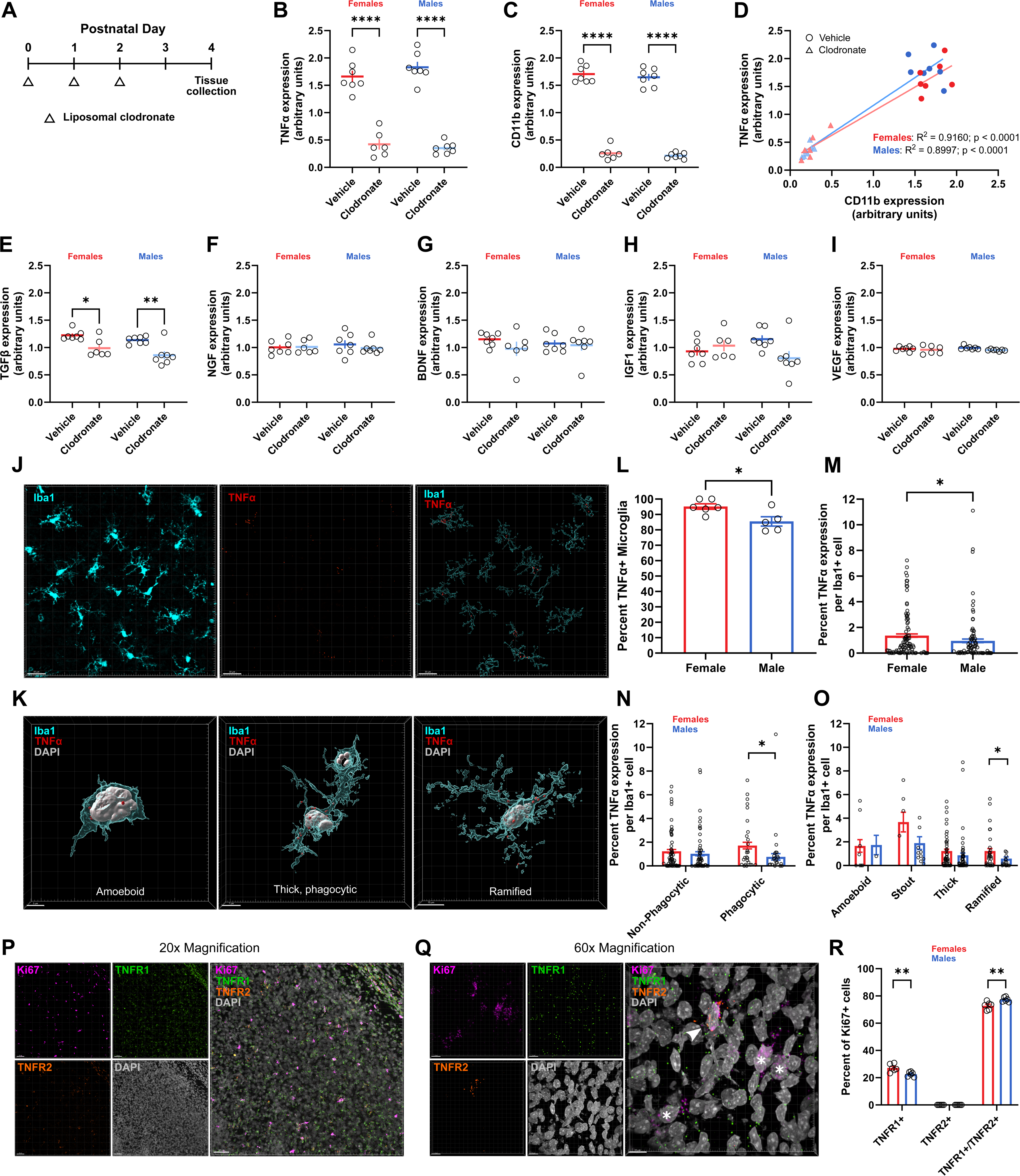
Expression of Microglial TNFα and TNF Receptors are Sex-Specific in the Developing Amygdala. (A) Schematic showing the experimental treatments and timeline for (B) – (I). (B) Quantification of TNFα expression in whole amygdala tissue by qPCR. Two-way ANOVA main effect of treatment F(1, 23) = 230.9; p < 0.0001. Bonferroni’s post hoc comparisons are shown, n = 6-7 rats per group. (C) Quantification of CD11b expression in whole amygdala tissue by qPCR. Two-way ANOVA main effect of treatment F(1, 23) = 790.4; p < 0.0001. Bonferroni’s post hoc comparisons are shown, n = 6-7 rats per group. (D) Correlation between TNFα and CD11b expression (from B and C). Points indicate values for individual animals; colors indicate sex while shape indicates treatment (circles = vehicle, diamonds = clodronate). Blue and red lines indicate linear regression for males and females, respectively. (E) Quantification of TGFβ expression in whole amygdala tissue by qPCR. Two-way ANOVA main effect of treatment F(1, 23) = 22.67; p < 0.0001. Bonferroni’s post hoc comparisons are shown, n = 6-7 rats per group. (F) Quantification of NGF expression in whole amygdala tissue by qPCR. n = 6-7 rats per group. (G) Quantification of BDNF expression in whole amygdala tissue by qPCR. n = 6-7 rats per group. (H) Quantification of IGF1 expression in whole amygdala tissue by qPCR. n = 6-7 rats per group. (I) Quantification of VEGF expression in whole amygdala tissue by qPCR. n = 6-7 rats per group. (J) Representative maximum intensity projection of the amygdala immunolabeled for Iba1 (top left) and RNAscope signal for TNFα (bottom left). Three-dimensional rendering (right) shows colocalization of TNFα transcripts with Iba1 signal. Scale bars represent 30 µm. (K) Three-dimensional reconstructions of amoeboid (left), thick, phagocytic (middle), and ramified (right) microglia morphologies showing representative TNFα transcript colocalization. Scale bars represent 5 µm (left), 7 µm (middle), and 10 µm (right). (L) Quantification of the percentage of microglia that are TNFα+. Student’s t test t(9) = 2.845; p = 0.0192. n = 5 males (150 cells) and 6 females (187 cells). (M) Quantification of TNFα transcript colocalization within male and female microglia in the amygdala. Student’s t test t(335) = 2.148; p = 0.0325, n = 5 males (150 cells) and 6 females (187 cells). (N) Quantification of TNFα transcript colocalization within male and female microglia (from G) separated by phagocytic status. Phagocytic microglia: student’s t test t(91) = 2.423; p = 0.0174, n = 5 males (42 cells) and 6 females (51 cells). (O) Quantification of TNFα transcript colocalization within male and female microglia (from G) separated by morphology. Ramified microglia: student’s t test t(93) = 2.110; p = 0.0375, n = 5 males (34 cells) and 6 females (61 cells). (P) Representative maximum intensity projection of the amygdala showing RNAscope signal for Ki67 (top left), TNFR2 (bottom left), TNFR1 (top middle), DAPI (bottom middle) and the resulting merged image (right). Scale bars represent 50 µm. (Q) Representative maximum intensity projection of the amygdala showing RNAscope signal for Ki67 (top left), TNFR2 (bottom left), TNFR1 (top middle), DAPI (bottom middle) and the resulting merged image (right). Scale bars represent 10 µm. White arrowheads indicates Ki67+/TNFR1+/TNFR2+ cell, white asterisks indicate Ki67+/TNFR1+ cells. (R) Quantification of the percentage of Ki67+ cells colocalizing with TNFR1 and TNFR2 transcripts. TNFR1+: student’s t test t(10) = 3.313; p = 0.0078. TNFR1+/TNFR2+: student’s t test t(10) = 3.313; p = 0.0078; n = 6 rats per group. Bars represent the mean ± SEM. Open circles represent individual data points for each animal. *p < 0.05, **p < 0.01, ***p < 0.001, ****p < 0.0001.

To determine whether *TNFα* expression was related to a particular microglial phenotype or morphology, we combined immunohistochemistry for Iba1 together with RNAscope *in situ* hybridization for *TNFα* and quantified *TNFα* expression within individual microglia in the male and female amygdala at P4 (Figure 3J, 3K). Overall, the vast majority of microglia were *TNFα*+; however, females had a significantly higher proportion of *TNFα*+ microglia than males (95.17% in females vs 85.51% in males; Figure 3L). Moreover, the volume of *TNFα* colocalization was higher in female microglia than male (1.366% in females vs 0.952% in males; Figure 3M), and the difference in colocalization was greatest when comparing phagocytic microglia (1.714% in females vs 0.763% in males; Figure 3N). When looking across the four “historical” morphological categories of microglia in development (amoeboid, stout, thick, or ramified; Schwarz et al., 2012), we found similar *TNFα* expression in all types of microglia except ramified, where female ramified microglia again had greater *TNFa* colocalization compared to male (1.225% in females vs 0.583% in males; Figure 3O).

We then used RNAscope to determine the localization patterns of the TNFα receptors, *TNFR1* and *TNFR2*, on newborn cells (identified as *Ki67*+) in the developing amygdala (Figure 3P, 3Q). Approximately 25% of *Ki67*+ cells colocalized with only *TNFR1*, with this percentage being higher in females compared to males (27.19% in females vs 22.63% in males). Conversely, nearly 75% of *Ki67*+ cells colocalized with both *TNFR1* and *TNFR2*, with the percentage being higher in males compared to females (77.37% in males vs 72.81% in females). In our analysis, no *Ki67*+ cells colocalized with only *TNFR2* in either males or females (Figure 3R). Together, these data show that microglia are the primary source of TNFα in the developing amygdala, and that TNF receptor expression is enriched on newborn cells.

### Inhibition of TNFα-induced NF-κB Activation Reveals Female-Biased Cell Survival in the Developing Amygdala and Programs Juvenile Play

To test the hypothesis that microglial TNFα signaling in females was responsible for the observed higher rates of cell survival, we used the anti-inflammatory compound celastrol, which inhibits TNFα-induced NF-κB activation (Lee et al., 2006; Jung et al., 2007; Sethi et al., 2007). We hypothesized that blocking the downstream signaling of TNFα in the brain would effectively mimic the effects of microglia depletion (i.e. decreased TNFα expression) on brain development and behavior. Similar to the previous experimental paradigm used to investigate cell survival (as in Figure 2A), we treated male and female pups with BrdU on P0 and celastrol intra-amygdala infusions from P1-3, then quantified BrdU+ cells on P4 (Figure 4A). As predicted, celastrol treatment in females significantly reduced BrdU+ cell number to male levels, while BrdU+ cell number was not affected in males (Figure 4B).

**Figure 4.**
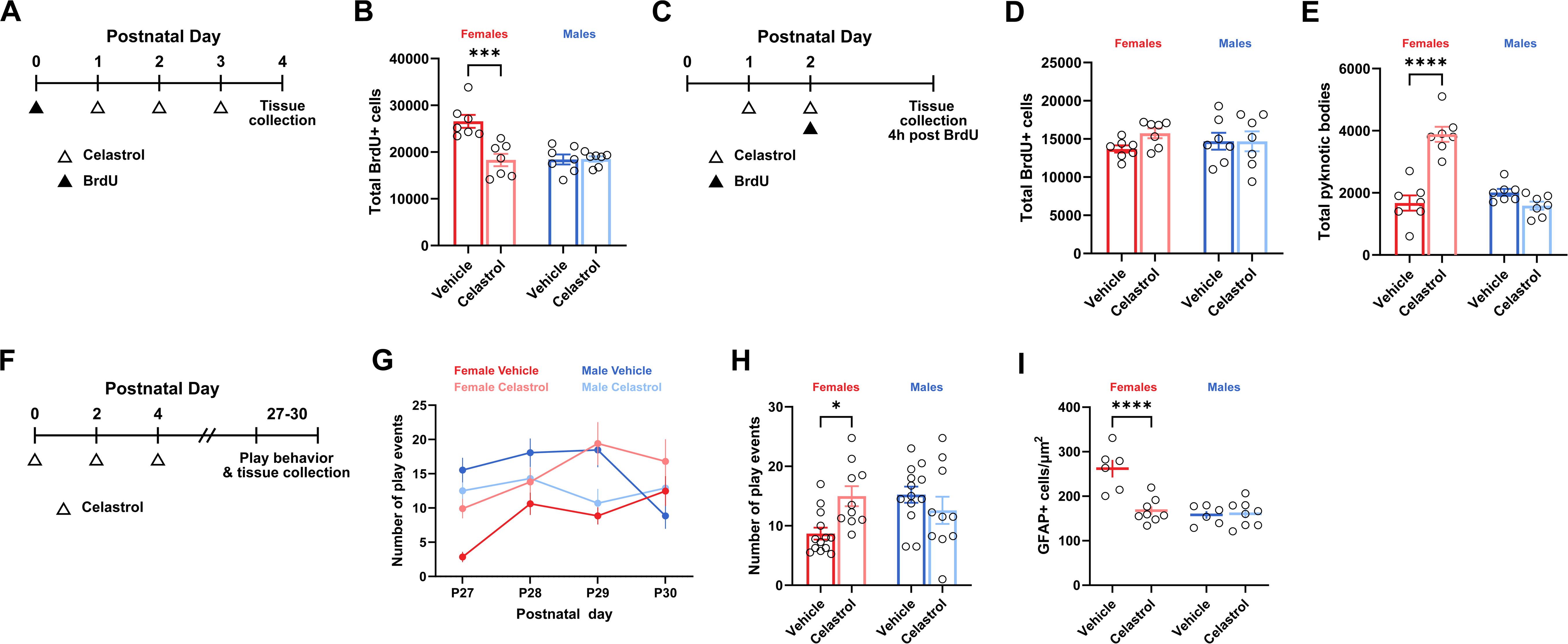
NF-κB Inhibition Reduces Newborn Cell Survival and Increases Play Behavior in Females, but Not Males. (A) Schematic showing the experimental treatments and timeline for (B). (B) Quantification of the number of BrdU+ cells. Two-way ANOVA sex x treatment interaction F(1, 24) = 13.69; p = 0.0011. Bonferroni’s post hoc comparisons are shown, n = 7 rats per group. (C) Schematic showing the experimental treatments and timeline for (D) – (E). (D) Quantification of the number of BrdU+ cells. n = 7 rats per group. (E) Quantification of the number of pyknotic bodies. Two-way ANOVA sex x treatment interaction F(1, 24) = 47.01; p < 0.0001. Bonferroni’s post hoc comparisons are shown, n = 7 rats per group. (F) Schematic showing the experimental treatments and timeline for (G) – (I). (G) Quantification of the number of play events daily from P27 – P30. Colors indicate sex and treatment. (H) Quantification of the mean number of play events from P27 – P30. Two-way ANOVA sex x treatment interaction F(1, 42) = 8.052; p = 0.0070. Bonferroni’s post hoc comparisons shown, n = 10-13 rats per group. (I) Quantification of the density of GFAP+ cells in the MePD. Two-way ANOVA sex x treatment interaction F(1, 24) = 15.68; p = 0.0006. Bonferroni’s post hoc comparisons shown, n = 6-8 rats per group. Bars and line graphs represent the mean ± SEM. Open circles represent individual data points for each animal. MePD, posterodorsal medial amygdala. *p < 0.05, **p < 0.01, ***p < 0.001, ****p < 0.0001.

In order to determine whether NF-κB inhibition also altered cell proliferation and cell death, we treated male and female pups with celastrol on P1 and P2, and BrdU on P2 followed by tissue collection 4 hours later (Figure 4C). Celastrol treatment did not affect BrdU+ cell number in males or females (Figure 4D). However, celastrol treatment selectively increased the number of pyknotic bodies in females compared to controls (Figure 4E). This is consistent with TNFa mediating cell survival, not cell proliferation.

Finally, we treated male and female pups with celastrol on P0, P2, and P4 and quantified play behavior beginning on P27 to determine the impact of neonatal NF-κB inhibition on later-life behavior and astrocyte density in the MePD (Figure 4F). As predicted, celastrol-treated females were behaviorally masculinized and played significantly more than vehicle-treated control females, while males were unaffected by celastrol treatment and continued to play at levels significantly higher than control females (Figure 4G, 4H). Celastrol treatment also significantly reduced GFAP+ cell density only in females, while males were again unaffected (Figure 4I). Together, these data demonstrate that TNFα induction of NF-κB regulates newborn astrocyte survival selectively in the developing female amygdala to feminize later-life juvenile play behavior.

## DISCUSSION

Juvenile rough-and-tumble play is a unique social behavior-it is expressed only briefly during early life, occurs independent of reproductive competency or aggression, and has no immediate impact on reproductive fitness (Burghardt et al., 2024). Yet a pan-species feature of play is the consistently higher frequency and intensity of play by males compared to females (VanRyzin et al., 2020b). Recent studies have revealed that juvenile play imparts later-life reproductive fitness benefits in males (Marquardt et al., 2023; Holmes et al., 2024), which could explain the tightly regulated control of masculinization of neural circuitry underlying play (Pellis 2002; Bredewold et al., 2014; Dumais & Veenema, 2016; VanRyzin et al., 2019). What remains a mystery, however, is whether play confers any benefits to females, or if the expression of play in females is simply an inconsequential evolutionary byproduct. The data we present here argue against the latter hypothesis and demonstrate that the feminization of play is as tightly regulated and exquisitely controlled as masculinization (Figure 5). We find that while microglia regulate basal cell proliferation in the developing amygdala in both sexes, microglia in the developing female brain actively support the survival of these newborn cells, primarily astrocytes, through production of TNFα and activation of NF-κB signaling. Increased numbers of astrocytes in the MePD is correlated with lower neuronal activation and an associated reduction in playfulness during the juvenile period (VanRyzin et al., 2019). These findings provide one of the first examples of active feminization of a brain region and associated behavioral consequences.

**Figure 5:**
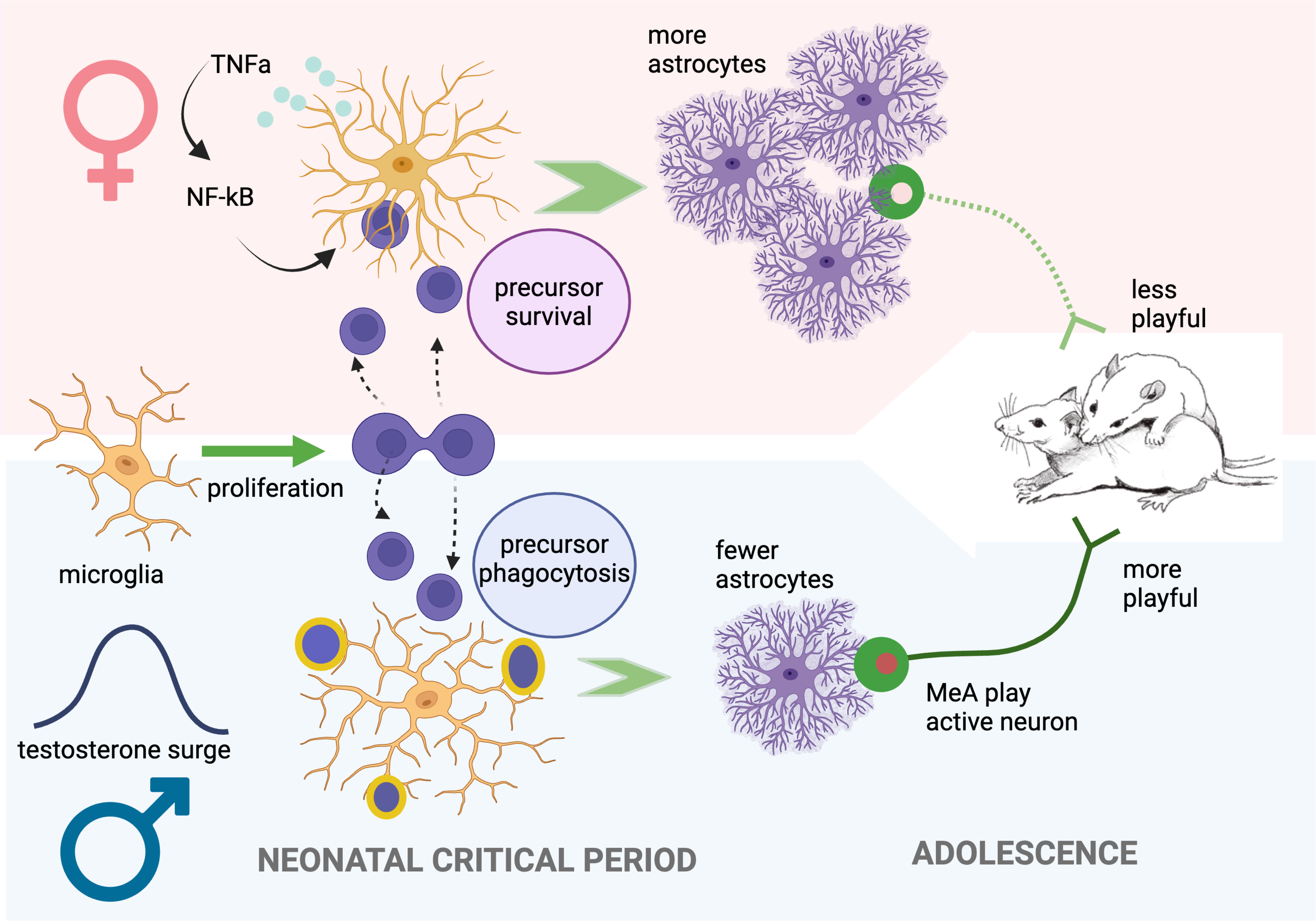
Schematic Summary of Feminization and Masculinization of Play Behavior. Microglia in the developing amygdala support the proliferation of newborn cells—mostly astrocytic precursors—in both sexes. In females, microglia play an additional role by promoting the survival of astrocytic precursors through a TNFa-NF-κB pathway. The pro-survival capacity of microglia is not found in male medial amygdala but there are high rates of phagocytosis of astrocyte precursors (VanRyzin et al., 2019). Both the loss of the pro-survival capacity and gain of phagocytic activity are induced by the androgen surge specific to males. As a result of these dual processes, adolescent females have significantly more mature astrocytes in the medial amygdala compared to males and this is associated with dampened neuronal activation, as evidenced by Erg1 expression (VanRyzin et al., 2019) during a play encounter, resulting in lower levels of play compared to males.

### Astrocyte density in the MePD is regulated by a balance of microglia functions during development

We previously demonstrated that microglia-mediated phagocytosis reduces the number of astrocytic precursors in the male medial amygdala (VanRyzin et al., 2019). Our findings here highlight an additional mechanism by which microglia modulate astrocyte numbers and do so differently in males and females: by supporting cell survival. These findings complement previous research showing that newborn cell number in the developing male amygdala is controlled by microglia phagocytosis (VanRyzin et al., 2019), and highlight dual opposing processes that shape sex differences in astrocyte number throughout development. To this point, our data show that the feminization of cell survival can be prevented by masculinizing female pups with testosterone during this developmental window, presumably through androgen-dependent induction of microglia phagocytosis (VanRyzin et al., 2019) and inhibition of the pro-survival effect of microglia. In this way, the architecture of the amygdala, and subsequent later-life play behavior, is the result of a balance between these microglia masculinization/feminization processes.

Interestingly, males and females were equally sensitive to the loss of microglia, as BrdU+ cell number was reduced when newborn cells were labeled after microglia depletion, suggesting that in the absence of microglia, cell proliferation within the amygdala decreases. This raises an interesting question: how are microglia increasing cell proliferation during development? Microglia are intricately intertwined with their microenvironments (Mosher et al., 2012; Ribeiro Xavier et al., 2015; Marsters et al., 2020; Rosin et al., 2021) and produce several growth factors and cytokines that support cell proliferation (Elkabes et al., 1996; Nicholas et al., 2001; Butovsky et al., 2006; Nakanishi et al., 2007; Ueno et al., 2013; Shigemoto-Mogami et al., 2014; Hagemeyer et al., 2017; Kreisel et al., 2019). Future studies are therefore needed to determine precisely which microglial factors underlie basal proliferation within the amygdala and whether the mechanisms to promote proliferation are the same in males and females.

### Newborn astrocyte survival is regulated by microglia-derived TNFα in the developing female amygdala

Our findings highlight TNFα as a key survival factor for newborn astrocytes, which serves as an agent of sexual differentiation of the amygdala. Similarly, TNFα directs sexual differentiation of the anteroventral periventricular (AVPV) nucleus of the hypothalamus by promoting neuron survival in females, resulting in a larger AVPV compared to males (Krishnan et al., 2009; Petersen et al., 2012). In the AVPV, the source of TNFα is unknown, but has been suggested to be of microglial origin (Nelson & Lenz, 2017; VanRyzin et al., 2018). Here, we show that microglia are the predominant producers of TNFα in the neonatal amygdala and that TNFα expression is higher in female microglia than male microglia. TNFα expression was not related to traditional morphological categories of microglia “activation” (Paolicelli et al., 2022), suggesting that basal TNFα likely does not act as a pro-inflammatory cytokine in development. Rather, TNFα expression was observed in microglia across the spectrum of morphologies.

We hypothesized that TNFa was acting directly on newborn cells to promote their survival. TNFα binds to either TNF receptor 1 (TNFR1) or receptor 2 (TNFR2) and activation of either receptor can lead to downstream activation of NF-κB and cell survival, despite the presence of a “death domain” on the intracellular tail of TNFR1 (Locksley et al., 2001; Probert, 2015; Gonzalez Caldito, 2023). While TNFR1 is ubiquitously expressed by many cells in the brain, TNFR2 expression is largely restricted to immune cells, oligodendrocytes, and subsets of neurons in the brain (Probert, 2015). We found that TNFR1 and TNFR2 are expressed together on the majority (∼75%) of newborn cells, with the remainder of newborn cells expressing only TNFR1. Whether this differential expression has any biological significance for amygdala development is unknown; it may be that co-expression of TNFR1 and TNFR2 is related to progenitor subtype, as over 80% of newborn cells in the post-natal amygdala differentiate into astrocytes (VanRyzin et al., 2019). Furthermore, our studies implicate TNFα acting on TNFR1 and TNFR2 as regulating newborn cell survival, consistent with others (Arnett et al., 2001; Shigemoto-Mogami et al., 2014; Nelson et al., 2020), but do not determine whether one, or both, receptors are necessary for this effect. Additional studies are needed to more fully elucidate these signaling mechanisms, as each receptor is coupled to distinct cellular signaling cascades that converge upon NF-κB activation and lead to cell survival (Ting & Bertrand, 2017; Medler et al., 2022)

We further hypothesized that the survival effects of TNFa were mediated by NF-kB. To that end, we used pharmacological inhibition of TNFα-induced NF-κB activation and found it sufficient to decrease newborn cell survival in the neonatal amygdala. NF-κB signaling is a convergence point for cell survival, development, and maturation in the brain (Nicholas et al., 2001; Methot et al., 2013; 2018; Yamanishi et al., 2015), and can be activated by numerous cytokines and growth factors (Guo et al., 2024). Of the factors we assessed, TNFα was most affected by microglia depletion and despite well-documented roles for IGF1 (Beck et al., 1995; O’Kusky et al., 2000; Pérez-Martín et al., 2010; Ueno et al., 2013), TGFβ (Carvalho Alcantara Gomes et al., 2005; Meyers & Kessler, 2017), NGF (Friedman & Greene, 1999; Sanes et al., 2011), BDNF (Kimoto et al., 2009), and VEGF (Nowaka & Obuchowicz et al., 2012; Lelli et al., 2013; Kreisel et al., 2019; Okabe et al., 2020), the expression of each of these transcripts was largely unchanged in the neonatal amygdala following microglia depletion. This does not, however, rule out the possibility that each may have distinct and important roles in amygdala development.

These findings confirm the importance of astrocyte density to regulating the lower level of playfulness in females as there was an increase in playfulness observed when microglia were temporarily depleted or activation of the NF-κB pathway was temporarily blocked, with both inducing enduring impacts on astrocyte density. This suggests that active feminization governs the amount of playfulness in females. However, whether there is evolutionary benefit to females to be less playful, versus a detriment to females for being too playful, or whether play behavior is coincidentally co-regulated together with some other ancillary role for astrocyte density, remain open questions.

## Conclusion

In conclusion, we demonstrate a novel mechanism underlying feminization of the developing amygdala and female-typical juvenile play. Microglia are central to this process, as microglial derived TNFα supports newborn cell survival selectively in females which ultimately increases astrocyte density within the medial amygdala.

Uncovering mechanisms by which sex differences arise in development leads to new insights into natural sources of individual variation in brain development. In recent years, microglia have been found to be essential to driving sex differences in numerous brain regions through their many functions to influence the development of sex-specific behaviors (Lenz et al., 2013; Kopec et al., 2018; VanRyzin et al., 2019; Pickett et al., 2023). This is particularly salient for the medial amygdala, which is essential for processing social information in a sex-specific manner and regulating male aggression, female mating, and parental care behaviors (Choi et al., 2005; Bergan et al., 2014; Unger et al., 2015; Ishii et al., 2017; Li et al., 2017; Lischinsky et al., 2017). Together, our findings highlight that microglia—in the same brain region and at the same time in development—may carry out multiple functions which differ by sex, and shed light onto new ways by which brain development may go awry.

## Acknowledgements

This work was funded by National Institutes of Health grants R01DA039062 to MMM and F31MH123025 to AEM. We thank the University of Maryland School of Medicine’s Confocal Microscopy Core (Baltimore, Maryland), including J. Mauban, for technical assistance.

## REFERENCES

1. Ashwell K.W.S., Paxinos G. 2008. Atlas of the developing rat nervous system. Third edition (Academic Press).

2. Argue K.J., VanRyzin J.W., Falvo D.J., Whitaker A.R., Yu S.J., McCarthy M.M. 2017. Activation of both CB1 and CB2 endocannabinoid receptors is critical for masculinization of the developing amygdala and juvenile social play behavior. eNeuro 4, ENEURO.0344-16.2017.

3. Arnett H.A., Mason J., Marino M., Suzuki K., Matsushima G.K., Ting J.P. 2001. TNF alpha promotes proliferation of oligodendrocyte progenitors and remyelination. Nat Neurosci 4, 1116–1122.

4. Beck K.D., Powell-Braxton L., Widmer H.R., Valverde J., Hefti F. 1995. IGF1 gene disruption results in reduced brain size, CNO hypomyelination, and loss of hippocampal granule and striatal parvalbumin-containing neurons. Neuron 14, 717–730.

5. Bergan J.F., Ben-Shaul Y., Dulac C. 2014. Sex-specific processing of social cues in the medial amygdala. Elife 3, e02743.

6. Bredewold R., Smith C.J.W., Dumais K.M., Veenema A.H. 2014. Sex-specific modulation of juvenile social play behavior by vasopressin and oxytocin depends on social context. Front Behav Neurosci 8, 216.

7. Burghardt G.M., Pellis S.M., Schank J.C., Smaldino P.E., Vanderschuren L.J.M.J., Palagi E. 2024. Animal play and evolution: Seven timely research issues about enigmatic phenomena. Neurosci Biobehav Rev 160, 105617.

8. Butovsky O., Ziv Y., Schwartz A., Landa G., Talpalar A.E., Pluchino S., Martino G., Schwartz M. 2006. Microglia activated by IL-4 or IFN-γ differentially induce neurogenesis and oligodendrogenesis from adult stem/progenitor cells. Mol Cell Neurosci 31, 149–160.

9. Carvalho Alcantara Gomes F., de Oliveira Sousa V., Romão L. 2005. Emerging roles for TGF-beta1 in nervous system development. Int J Dev Neurosci 23, 413–424.

10. Choi G.B., Dong H-W., Murphy A.J., Valenzuela D.M., Yancopoulos G.D., Swanson L.W., Anderson D.J. 2005. Lhx6 delineates a pathway mediating innate reproductive behaviors from the amygdala to the hypothalamus. Neuron 46, 647–60.

11. Cunningham C.L., Martinez-Cerdeño V., Noctor S.C. 2013. Microglia regulate the number of neural precursor cells in the developing cerebral cortex. J Neurosci 33, 4216–4233.

12. Dumais K.M., Veenema A.H. 2016. Vasopressin and oxytocin receptor systems in the brain: Sex differences and sex-specific regulation of social behavior. Front Neuroendocrinol 40, 1–23.

13. Elkabes S., DiCicco-Bloom E.M., Black I.B. 1996. Brain microglia/macrophages express neurotrophins that selectively regulate microglia proliferation and function. J Neurosci 16, 2508–1521.

14. Forger N.G. 2016. Epigenetic mechanisms in sexual differentiation of the brain and behaviour. Philos Trans R Soc Lond B Biol Sci 371, 20150114.

15. Friedman W.J., Greene L.A. 1999. Neurotrophin signaling via Trks and p75. Exp Cell Res 253, 131–142.

16. Gonzalez Caldito N. 2023. Role of tumor necrosis factor-alpha in the central nervous system: a focus on autoimmune disorders. Front Immunol 14, 1213448.

17. Goodson J.L. 2005. The vertebrate social behavior network: evolutionary themes and variations. Horm Behav 48, 11–22.

18. Guo Q., Jin Y., Chen X., Ye X., Shen X., Lin M., Zeng C., Zhou T., Zhang J. 2024. NF-κB in biology and targeted therapy: new insights and translational implications. Signal Transduct Target Ther 9, 53.

19. Hagemeyer N., Hanft K.M., Akriditou M.A., Unger N., Park E.S., Stanely E.R., Staszewski O., Dimou L., Prinz M. 2017. Microglia contribute to normal myelinogenesis and to olidogendrocyte progenitor maintenance during adulthood. Acta Neuropathol 134, 441–458.

20. Holmes K.G., Krützen M., Ridley A.R., Allen S.J., Connor R.C., Gerber L., Flaherty Stamm C., King S.L. 2024. Juvenile social play predicts adult reproductive success in male bottlenose dolphins. Proc Natl Acad Sci U S A 121, e2305948121.

21. Ishii K.K., Osakada T., Mori H., Miyasak N., Yoshihara Y., Miyamichi K., Touhara K. 2017. A labeled-line neural circuit for pheromone-mediated sexual behaviors in mice. Neuron 95, 123–137.e8.

22. Ji K., Akgul G., Wollmuth L.P., Tsirka S.E. 2013. Microglia actively regulate the number of functional synapses. PLoS ONE 8, e56293.

23. Jung H.W., Chung Y.S., Kim Y. S., Park Y-K. 2007. Celastrol inhibits production of nitric oxide and proinflammatory cytokines through MAPK signal transduction and NF-kappaB in LPS-stimulated BV-2 microglial cells. Exp Mol Med 39, 715–721.

24. Kimoto H., Eto R., Abe M., Kato H., Araki T. 2009. Alterations of glial cells in the mouse hippocampus during postnatal development. Cell Mol Neurobiol 29, 1181–1189.

25. Kopec A.M., Smith C.J., Ayre N.R., Sweat S.C., Bilbo S.D. 2018. Microglial dopamine receptor elimination defines sex-specific nucleus accumbens development and social behavior in adolescent rats. Nat Commun 9, 3769.

26. Krebs-Kraft D.L., Hill M.N., Hillard C.J., McCarthy M.M. 2010. Sex difference in cell proliferation in developing rat amygdala mediated by endocannabinoids has implications for social behavior. Proc. Natl. Acad. Sci. USA 107, 20535–20540.

27. Kreisel T., Wolf B., Keshet E., Licht T. 2019. Unique role for dentate gyrus microglia in neuroblast survival and in VEGF-induced activation. Glia 67, 594–619.

28. Krishnan S., Intlekofer K. A., Aggison L.K., Petersen S.L. 2009. Central role of TRAF-interacting protein in a new model of brain sexual differentiation. Proc. Natl. Acad. Sci. USA 106(32), 16692–16697.

29. Lee J.H., Koo T.H., Yoon H., Jung H.S., Jin H.Z., Lee K., Hong Y.S., Lee J.J. 2006. Inhibition of NF-kappaB activation through targeting I kappa B kinase by celastrol, a quinone method triterpenoid. Biochem Pharmacol 72, 1311–1321.

30. Lenz K.M., Nugent B.M., Haliyur R., McCarthy M.M. 2013. Microglia are essential to masculinization of brain and behavior. J Neurosci 33(7), 2761–2772.

31. Lelli A., Gervais A., Colin C., Chéret C., de Almodovar C.R., Carmeliet P., Krause K-H., Boilée S., Mallat M. 2013. The NADPH oxidase Nox2 regulates VEGFR1/CSF-1R-mediated microglial chemotaxis and promotes early postnatal infiltration of phagocytes in the subventricular zone of the mouse cerebral cortex. Glia 61, 1542–1555.

32. Li Y., Mathis A., Grewe B.F., Osterhout J.A., Ahanonu B., Schnitzer M.J., Murthy V.N., Dulac C. 2017. Neuronal representation of social information in the medial amygdala of awake behaving mice. Cell 171, 1176–1190.e17.

33. Lischinsky J.E., Sokolowski K., Li P., Esumi S., Kamal Y., Goodrich M., Oboti L., Hammond T.R., Krishnamoorthy M., Feldman D., Huntsman M., Liu J., Corbin J.G. 2017. Embryonic transcription factor expression in mice predicts medial amygdala neuronal identity and sex-specific repsponses to innate behavioral cues. Elife 6, e21012.

34. Lischinsky J.E., Yin L., Shi C., Prakash N., Burke J., Shekaran G., Grba M., Corbin J.G., Lin D. 2023. Transcriptionally defined amygdala subpopulations play distinct roles in innate social behaviors. Nat Neurosci 26, 2131–2146.

35. Locksley R.M., Killeen N., Lenardo M.J. 2001. The TNF and TNF receptor superfamilies: integrating mammalian biology. Cell 104, 487–501.

36. Marin-Teva J.L., Dusart J., Colin C., Gervais A., van Rooijen N., Mallat M. 2004. Microglia promote the death of developing Purkinje cells. Neuron 41, 535–547.

37. Marquardt A.E., VanRyzin J.W., Fuquen R.W., McCarthy M.M. 2023. Social play experience in juvenile rats is indispensable for appropriate socio-sexual behavior in adulthood in males but not females. Front Behav Neurosci 16, 1076765.

38. Marsters C.M., Nesan D., Far R., Klenin N., Pittman Q.J., Kurrasch D.M. 2020. Embryonic microglia influence developing hypothalamic glial populations. J Neuroinflammation 17, 146.

39. McCarthy M.M., Arnold A.P. 2011. Reframing sexual differentiation of the brain. Nat Neurosci 14, 677–683.

40. McCarthy M.M., de Vries G.J., Forger N.G. 2017. Sexual differentiation of the brain: a fresh look at mode, mechanisms, and meaning, in: Pfaff DW and Joels M. (eds), Hormones, Brain and Behavior. Academic Press, Boston, pp. 3–32.

41. Meaney M.J., Stewart J., Poulin P., McEwen B.S. 1983. Sexual differentiation of social play in rat pups is mediated by the neonatal androgen-receptor system. Neuroendocrinology 37, 85–90.

42. Meaney M.J., McEwen B.S. 1986. Testosterone implants into the amygdala during the neonatal period masculinize the social play of juvenile female rats. Brain Res. 398, 324–238.

43. Medler J., Kucka K., Wajant H. 2022. Tumor necrosis factor receptor 2 (TNFR2): An emerging target in cancer therapy. Cancers, 14, 2603.

44. Methot L., Hermann R., Tang Y., Lo R., Al-Jehani H., Jhas S., Svoboda D., Slack R.S., Barker P.A., Stifani S. 2013. Interaction and antagonistic roles of NF-κB and Hes6 in the regulation of cortical neurogenesis. Mol Cell Biol 33, 2797–2808.

45. Methot L., Soubannier V., Hermann R., Campos E., Li S., Stifani S. 2018. Nuclear factor-kappaB regulates multiple steps of gliogenesis in the developing murine cerebral cortex. Glia 66, 2659–2672.

46. Meyers E.A., Kessler J.A. 2017. TGF-β family signaling in neural and neuronal differentiation, development, and function. Cold Spring Harb Perspect Biol 9, a022244.

47. Miyamoto A., Wake H., Ishikawa A.W., Eto K., Shibata K., Murakoshi H., Koizumi S., Moorhouse A.J., Yoshimura Y., Nabekura J. 2016. Microglia contact induces synapse formation in developing somatosensory cortex. Nat Commun 7, 12540.

48. Mosher K.I., Andres R.H., Fukuhara T., Bieri G., Hasegawa-Moriyama M., Yingbo H., Guzman R., Wyss-Coray T. 2012. Neural progenitor cells regulate microglia functions and activity. Nat Neurosci 15, 1485–1487.

49. Nakanishi M., Niidome T., Matsuda S., Akaike A., Kihara T., Sugimoto H. 2007. Microglia-derived interleukin-6 and leukaemia inhibitory factor promote astrocytic differentiation of neural stem/progenitor cells. Eur J Neurosci 25, 649–658.

50. Nelson L.H., Lenz K.M. 2017. The immune system as a novel regulator of sex differences in brain and behavioral development. J Neurosci Res 95, 447–461.

51. Nelson L.H., Peketi P., Lenz K.M. 2021. Microglia regulate cell genesis in a sex-dependent manner in the neonatal hippocampus. Neuroscience 453, 237–255.

52. Newman S.W. 1999. The medial extended amygdala in male reproductive behavior: a node in the mammalian social behavior network. Ann N Y Acad Sci 877, 242–257.

53. Nicholas R.S.T.J., Wing M.G., Compston A. 2001. Nonactivated microglia promote oligodendrocyte precursor survival and maturation through the transcription factor NF-κB. Eur J Neurosci 13, 959–967.

54. Nowaka M.M., Obuchowicz E. 2012. Vascular endothelial growth factor (VEGF) and its role in the central nervous system: a new element in the neurotrophic hypothesis of antidepressant drug action. Neuropeptides 46, 1–10.

55. Okabe K., Fukada H.., Tai-Nagara I., Ando T., Honda T., Nakajima K., Takeda N., Fong G-H., Ema M., Kubota Y. 2020. Neuron-derived VEGF contributes to cortical and hippocampal development independently of VEGFR1/2-mediated neurotropism. Dev Biol 459, 65–71.

56. O’Kusky J.R., Ye P., D’Ercole A.J. 2000. Insulin-like growth factor 1 promotes neurogenesis and synaptogenesis in the hippocampal dentate gyrus during postnatal development. J Neurosci, 20, 8435–8442.

57. Paolicelli R.C., Bolasco G., Pagani F., Maggi L., Scianni M., Panzanelli P., Giustetto M., Ferreira T.A., Guiducci E., Dumas L., Ragozzino D., Gross C.T. 2011. Synaptic pruning by microglia is necessary for normal brain development. Science 333, 1456–1458.

58. Paolicelli R.C., Sierra A., Stevens B., Tremblay M-E., Aguzzi A., Ajami B., Amit I., Audinat E., Bechmann I., Bennett M., Bennet F., Bessis A., Biber K., Bilbo S., Blurton-Jones M., Boddeke E., Brites D., Brone B., Brown G.C., Butovsky O., Carson M.J., Castellano B., Colonna M., Cowley S.A., Cunningham C., Davalos D., De Jager P.H., de Strooper B., Denes A., Eggen B.J.L., Eyo U., Galea E., Garel S., Ginhoux F., Glass C.K., Gokce O., Gomez-Nicola D., Gonzalez B., Gordon S., Graeber M.B., Greenhalgh A.D., Gressens P., Greter M., Gutmann D.H., Haass C., Heneka M.T., Heppner F.L., Hong S., Hume D.A., Jung S., Kettenmann H., Kipnis J., Koyama R., Lemke G., Lynch M., Majewska A., Malcangio M., Malm T., Mancuso R., Masuda T., Matteoli M., McColl B.W., Miron V.E., Molofsky A.V., Monje M., Mracsko E., Nadjar A., Neher J.J., Neniskyte U., Neumann H., Noda M., Peng B., Peri F., Perry V.H., Popovich P.G., Pridans C., Priller J., Prinz M., Ragozzino D., Ransohoff R.M., Salter M.W., Schaefer A., Schafer D.P., Schwartz M., Simons M., Smith C.J., Streit W.J., Tay T.L., Tsai L-H., Verkhratsky A., von Bernhardi R., Wake H., Wattamer V., Wolf S.A., Wu L-J., Wyss-Coray T. 2022. Microglia states and nomenclature: A field at its crossroads. Neuron 110, 3458–3483.

59. Pellis S.M. 2002. Sex differences in play fighting revisited: traditional and non-traditional mechanisms of sexual differentiation in rats. Arch Sex Behav 31, 17–26.

60. Pérez-Martín M., Cifuentes M., Grondona J.M., López-Avalos M.D., Gómez-Pinedo U., García-Verdugo J.M., Fernández-Llebrez P. 2010. IGF-I stimulates neurogenesis in the hypothalamus of adult rats. Eur J Neursci 31, 1533–1548.

61. Petersen S.L., Krishnan S., Aggison L.K., Intlekofer K.A., Moura P.J. 2012. Sexual differentiation of the gonadotropin surge release mechanism: a new role for the canonical NFκB signaling pathway. Front. Neuroendocrinol. 33(1), 36–44.

62. Pickett L.A., VanRyzin J.W., Marquardt A.E., McCarthy M.M. 2023. Microglia phagocytosis mediates the volume and function of the sexually dimorphic nucleus of the preoptic area. Proc Natl Acad Sci U S A 120(10), e2212646120.

63. Ponti G., Obernier K., Guinto C., Jose L., Bonfanti L., Alvarez-Buylla A. 2013. Cell cycle and lineage progression of neural progenitors in the ventricular-subventricular zones of adult mice. Proc Natl Acad Sci U S A 110, E1045–1054.

64. Probert L. 2015. TNF and its receptors in the CNS: The essential, the desirable and the deleterious effects. Neuroscience 302, 2–22.

65. Raam T., Hong W. 2021. Organization of neural circuits underlying social behavior: A consideration of the medial amygdala. Curr Opin Neurobiol 68, 124–136.

66. Ribeiro Xavier A.L., Kress B.T., Goldman S.A., Lacerda de Menezes J.R., Nedergaard M.A. 2015. Distinct population of microglia supports adult neurogenesis in the subventricular zone. J Neurosci 35, 11848–11861.

67. Rosin J.M., Sinha S., Biernaskie J., Kurrasch D.M. 2021. A subpopulation of embryonic microglia respond to maternal stress and influence nearby neural progenitors. Dev Cell 56, 1326–1345.e6.

68. Sanes D.H., Thomas A.R., Harris W.A. 2011. Naturally-occurring cell death, in: Development of the Nervous System, Third Edition, Academic Press, Boston, pp. 171–208.

69. Schafer D.P., Lehrman E.K., Kautzman A.G., Koyama R., Mardinly A.R., Yamasaki R., Ransohoff R.M., Greenberg M.E., Barres B.A., Stevens B. 2012. Microglia sculpt postnatal neural circuits in an activity and complement-dependent manner. Neuron 74, 691–705.

70. Schmittgen T. D., Livak K.J. 2008. Analyzing real-time PCR data by the comparative C(T) method. Nat Protoc 3(6), 1101–1108.

71. Schwarz J.M., Sholar P.W., Bilbo S.D. 2012. Sex differences in microglial colonization of the developing rat brain. J. Neurochem. 120, 948–963.

72. Sethi G., Seok Ahn K., Pandey M.K., Aggarwal B.B. 2007. Celastrol, a novel triterpene, potentiates TNF-induced apoptosis and suppresses invasion of tumor cells by inhibiting NF-kappaB-regulated gene products and TAK1-mediated NF-kappaB activation. Blood 109, 2727–2735.

73. Shigemoto-Mogami Y., Hoshikawa K., Goldman J.E., Sekino Y., Sato K. 2014. Microglia enhance neurogenesis and oligodendrogenesis in the early postnatal subventricular zone. J Neurosci 34, 2231–2243.

74. Sierra A., Encinas J.M., Deudero J.J., Chancey J.H., Enikolopov G., Overstreet-Wadiche L.S., Tsirka S.E., Maletic-Savatic M. 2010. Microglia shape adult hippocampal neurogenesis through apoptosis-coupled phagocytosis. Cell Stem Cell 7, 483–495.

75. Squarzoni P., Oller G., Hoeffel G., Pont-Lezica L., Rostaing P., Low D., Bessis A., Ginhoux F., Garel S. 2014. Microglia modulate wiring of the embryonic forebrain. Cell Rep 8, 1271–1279.

76. Ting A.T., Bertrand M.J.M. 2017. More to life than NF-κB in TNFR1 signaling. Trends Immunol 37, 535–545.

77. Tsukahara S., Morishita M. 2020. Sexually dimorphic formation of the preoptic area and the bed nucleus of the stria terminalis by neuroestrogens. Front Neurosci 14, 797.

78. Ueno M., Fujita Y., Tanaka T., Nakamura Y., Kikuta J., Ishii M., Yamashita T. 2013. Layer V cortical neurons require microglial support for survival during postnatal development. Nat Neurosci 16, 543–551.

79. Unger E.K., Burke Jr K.H., Yang C.F., Bender K.J., Fuller P.M., Shah N.M. 2015. Medial amygdala aromatase neurons regulate aggression in both sexes. Cell Rep 10, 453–462.

80. VanRyzin J.W., Pickett L.A., McCarthy M.M. 2018. Microglia: Driving critical periods and sexual differentiation of the brain. Dev Neurobiol 78, 580–592.

81. VanRyzin J.W., Marquardt A.E., Argue K.J., Vecchiarelli H.A., Ashton S.E., Arambula S.E., Hill M.N., McCarthy M.M. 2019. Microglial phagocytosis of newborn cells is induced by endocannabinoids and sculpts sex differences in juvenile rat social play. Neuron 102(2), 435–449.e6.

82. VanRyzin J.W., Marquardt A.E., McCarthy M.M. 2020a. Assessing rough-and-tumble play behavior in juvenile rats. Bio Protoc 10(1), e3481.

83. VanRyzin J.W., Marquardt A.E., McCarthy M.M. 2020b. Developmental origins of sex differences in the neural circuitry of play. Int J Play 9, 58–75.

84. Yamanishi E., Yoon K., Alberi L., Galano N., Mizutani K. 2015. NF-κB signaling regulates the generation of intermediate progenitors in the developing neocortex. Genes Cells 20, 706–719.

85. Zuloaga D.G., Puts D.A., Jordan C.L., Breedlove S.M. 2008. The role of androgen receptors in the masculinization of brain and behavior: what we’ve learned from the testicular feminization mutation. Horm Behav 53, 613–626.

